# Therapeutic potential of Relaxin-2 in Heart Failure with preserved Ejection Fraction (HFpEF)

**DOI:** 10.64898/2026.05.14.725229

**Authors:** Joshua B. Palma, Bethann Gabris-Weber, Brenda McMahon, Anthony J. Mauro, Cynthia St. Hilaire, Rolando A. Cuevas, Thomas B. Dschietzig, Guillermo Romero, Guy Salama

## Abstract

**Aims:** Heart failure with preserved ejection fraction (HFpEF) afflicts millions annually and current treatments provide symptomatic relief. Here, we investigate the therapeutic potential of synthetic human Relaxin-2 (RLX) at reversing diastolic dysfunction (DD) and reducing arrhythmia vulnerability.

**Methods and Results:** Male ZSF1 rats were placed on a normal diet (ND, N=10 controls) or a high-fat diet (HFD, N=11), resulting in the development of DD in 11-weeks, based on serial echocardiograms (enlarged left atrium (LA), wall thickness, doppler flow: E/e’). Once HFpEF was confirmed, control and HFpEF rats were randomly treated with Relaxin (400µg/kg/day RLX, N=6) or the vehicle (N=5) for 2-weeks using implanted minipumps. Echocardiograms were repeated at weeks 1 and 2, then hearts were isolated, optically mapped, subjected to programmed electrical stimulation (PES) and tissues dissected for immuno-fluorescence (IF), and qPCR analysis. Circulating levels of glucose, RLX and NT-pro-ANP were measured, pre- and post-treatment. Echocardiograms indicated that RLX reversed DD by reducing LA dimensions and E/e’. Optical mapping revealed that 1/3 of HFpEF hearts exhibited sustained atrial and ventricular arrhythmia which were blocked by RLX as it tended to increase conduction velocity (CV). Based on IF, RLX increased Nav1.5, Connexin-43, β-catenin and Wnt1 expression. There were no significant changes in fibrosis in this HFpEF model. NT-pro-ANP was elevated in HFpEF and reduced towards control values by RLX. qPCR analysis showed that RLX decreased DKK1 and MMP1A and increased SCN5A expression compared to Vehicle treatment (N=6 and 5, respectively).

**Conclusions:** The ZSF1 model showed clear signs of HFpEF, including DD, enlargement of the LA, enhanced hemodynamic stress, increased vulnerability to sustained AF and VF, and elevated glucose and blood pressure. RLX treatment largely reversed DD, hemodynamic stress, and suppressed sustained arrhythmias. RLX elicited cardiac genomic changes, most likely through Wnt/canonical signaling, demonstrating RLX’s potential as a therapy for HFpEF.

## Introduction

The prevalence of heart failure (HF) has reached over 64 million people in the US, a devastating disease that is projected to increase with an aging population.(1) Among HF patients, ∼50% have HF with preserved ejection fraction (HFpEF), characterized by diastolic dysfunction while retaining an ejection fraction of over 50%.(1–3) Patients with HFpEF have a 6-9 times greater risk of supraventricular arrhythmias as a result of maladaptive remodeling of cardiac ion channels.(4, 5) Chronic hypertension and excess fibrosis in HFpEF lead to hypertrophy and contribute to the suppression of electrical coupling in atrial and ventricular syncytia.(6),(5) Atrial fibrillation often develops in HFpEF, synergistically with other phenotype changes, which exacerbates morbidity and mortality.(4–6) Patients with HFpEF are significantly older, more likely to be female, and more likely to have hypertension, obesity, anemia, atrial fibrillation, renal disease, and pulmonary disease compared to those with HFrEF (HF with reduced EF).(7) Current treatments consist of indirect approaches focused on the comorbidities of HFpEF. Given the abundant comorbidities associated with HFpEF, these treatments usually target symptoms of pulmonary edema/arterial hypertension, renal diseases, and metabolic syndrome/diabetes.(8)

Despite extensive basic science research and clinical trials, there are no treatments for HFpEF that significantly reverse its maladaptive cardiac remodeling.(9) More promising are recent treatments with sodium–glucose cotransporter 2 (SGLT2) inhibitors (∼ 2 years) which significantly reduced the risk of hospitalization for heart failure but left cardiovascular and all-cause mortality of HFpEF patients unchanged.(10, 11)

A major challenge in discovering effective treatments for HFpEF was the lack of suitable animal models that recapitulate many aspects of human disease. Genetic Models In (Indianapolis, IN) developed the ZSF1-obese rat by crossing male spontaneously hypertensive rats with female Zucker diabetic rats and it is generally accepted to closely mimic clinical HFpEF.(12), (13) ZSF1 rats placed on a high-fat diet (HFD) develop chronic hypertension, hyperlipidemia, glucose and exercise intolerance, and metabolic syndrome.(14) The interplay of these comorbidities leads to hypertrophy, arrhythmia, and arterial remodeling, while largely preserving systolic function.(14, 15) The manifestation of HFpEF in the ZSF1 model was verified by echocardiography through increased DD, filling pressures, and left atrial area.(14) The ZSF1-obese rat on a HFD results in a pathology close to human HFpEF and provides a unique opportunity to test for new therapeutic approaches.(13, 14, 16, 17)

Here, we tested synthetic human Relaxin-2 (RLX), an insulin-like hormone, as a possible therapy in the ZSF1 model of HFpEF. We previously showed that RLX reversed maladaptive remodeling in aged and chronically hypertensive rats and in pulmonary hypertension.(18–20) We found that RLX reversed excess collagen deposition and maladaptive fibrosis and markedly increased voltage-gated sodium channels at the level of mRNA, protein (Nav1.5) and current (I_Na_). RLX also increased connexin 43 (Cx43) expression and shifted Cx43 localization from the lateral membrane to the intercalated disks, and RLX reduced cellular hypertrophy.(19, 21, 22) The anti-arrhythmic properties of RLX were mediated by a reversal of maladaptive cardiovascular remodeling through genomic modifications of cardiac myocytes and fibroblasts by activating the Wnt1/β-catenin signaling pathway.(19) These actions of RLX suggest that this hormone might be an effective therapy for HFpEF since supraventricular arrhythmias are a common comorbidity in HFpEF patients.(4, 23) In this report, ZSF1-obese rats were placed on a HFD and serial echocardiograms were taken to track the development of HFpEF through changes in DD. Once HFpEF was diagnosed, minipumps were implanted and were randomly selected to receive RLX or the vehicle (controls) for two weeks, with echocardiograms taken at weeks 1 and 2 to track changes in HFpEF severity. At the end of the 2-week period, RLX or VEH treatment, the hearts were perfused in a Langendorff apparatus to optically map action potentials and Ca^2+^ transients and to evaluate changes in arrhythmia phenotype. Ventricular tissues were used for immunofluorescence (IF) and RT-PCR (Real Time-Polymerase Chain Reaction) to track changes in protein and gene expression, respectively and hence evaluate RLX’s therapeutic potential.

## Methods

### Animal Protocols

ZSF1-obese male rats (n=20), 9-weeks old were purchased from Charles River (Wilmington, MA; strain code: 378) and were acclimated for 1-week then randomly placed either on a standard Purina control diet (Purina ISO Pro Rodent 3000) (n=8 rats) or a HFD (Open-Source Diets; D12468 with 10 kcal% Soy Protein and 48 kcal% Lard) (n=12 rats) until week 22.

Of the 12 rats placed on the HFD, one rat died at week 14 and the remaining 11 rats developed HFpEF by week 18. At 20-weeks, the 11 rats which were on a HFD were implanted with osmotic minipumps (Alzet® Cupertino, CA; model 2ML2) and were randomly selected to receive RLX (400 µg/kg/day, n=6 HFpEF rats) or the vehicle (20 mmol/L Na^+^ acetate control; n=5 HFpEF rats, control treatment) for 14 days. The rats on a ND (n=8) were implanted with osmotic mini-pumps and randomly selected to receive RLX (400 µg/kg/day, n=4 normal rats) or the vehicle (20 mmol/L Na^+^ acetate control; n=4 normal rats) also for 14 days. The rats kept on the ND served as controls for the rats kept on the HFD.

The dose of RLX used in the current study is congruent with several studies in the literature. Previous reports on the effects of recombinant human RLX (rhRLX) on cardiac and renal fibrosis(24) and on arterial compliance(25) infused rhRLX in rats subcutaneously with osmotic mini-pumps at a dose of 0.5 mg/kg/day for 2 weeks. The latter dose resulted in ∼20-40 ng/ml of serum rhRLX (26), a level found on days 10-14 of gestation in pregnant rats.(27) Our studies on the anti-arrhythmic properties of recombinant human RLX in spontaneously hypertensive rats (SHR),(18) aged rats(19, 22, 28, 29) and rats with pulmonary arterial hypertension,(20) all used a slightly lower dose of 0.4 mg/kg/day also for 2 weeks. Preliminary studies on the effects of RLX on cardiac fibrosis indicated that 1 week of RLX treatment did not reverse fibrosis in SHR or aged (24 month) rats, whereas RLX treatments for 2 weeks effectively reversed fibrosis and was used as our standard protocol.(18, 22) In the current study, synthetic Human Relaxin-2 (shRLX) was obtained from Relaxera Pharmazeutische Gesellschaft mbH & Co. KG. shRLX is identical to human endogenous relaxin-2 in terms of structure and sequence.

### Experimental Protocol

Rats underwent phenotypic evaluation consisting of blood sample collections, weight measurements, and serial echocardiographic evaluations at weeks 10, 14, 18, 21 and 22. Ultrasound parameters used to evaluate HFpEF included: stroke volume, ejection fraction, and diastolic function through peak velocity of early (E) and late (A) mitral inflow signals and the ratio of E over e′ (peak velocity of early diastolic lateral mitral annular motion) which are indications of LV filling pressure. These parameters of HFpEF, measured at week 21 and 22 for rats that were treated with RLX were compared to those that were treated with vehicle. At 22 weeks, rats were evaluated by echocardiography under anesthesia then euthanized with a subcutaneous injection of Euthasol (96 mg/kg). Hearts were then isolated and perfused in a Langendorff apparatus for optical mapping of action potential with the voltage-sensitive dye, RH437 and free Ca^2+^ transients with the Ca^2+^-indicator, Rhod-2, as previously reported.(18, 30) After characterizing the arrhythmia phenotype, hearts were flash frozen, and ventricular tissues were sectioned for immuno-fluorescence or qPCR analysis. Studies were performed in accordance with the Guide for the Care and Use of Laboratory Animals and were approved by the Institutional Animal Care and Use Committee at the University of Pittsburgh.

### Echocardiograms

Ultrasound images were taken and analyzed at the University of Pittsburgh Small Animal Ultrasonography Core. Rats were anesthetized with 3% isoflurane and placed on a 37°C heat pad to maintain body temperature. During image acquisitions, isoflurane was maintained at 1–2% to maintain a heart rate between 250-400 beats per minute (bpm). Cardiac images were acquired with a Vevo 3100 imaging system and VisualSonic MX250 (15-30 MHz, 75μm axial resolution) linear array transducer (FUJIFILM, VisualSonics, Toronto, Canada). Systolic function was measured via the parasternal short-axis view at mid-papillary muscles. An M-Mode image was obtained to measure ejection fraction, fractional shortening, cardiac output, stroke volume, left ventricle (LV) Index, wall thickness, and internal diameters of the LV cavity. Mitral valve inflow was obtained by acquiring an apical 4-chamber view and placing a pulsed wave Doppler at the mitral valve tips to attain the E and A waves. The mitral valve tissue Doppler was obtained at the septal mitral valve annulus to obtain e’ and a’ measurements. The left atrial area was obtained in the apical 4-chamber view via tracing of the endocardial border. LV Index= LV mass (mg)/BSA(cm^2^) where BSA is the Body Surface Area in cm^2^ calculated from BSA = k X (rat weight in g)^2/3^ , where k=9.83 Meeh’s constant (k), as previously described by Gouma et al.,)(31) The Small Animal Sonography Core was supported by NIH grant number: 1S10OD023684-01A1

### Blood Pressure (BP) Measurements

The CODA BP System (Kent Scientific) was used to serially measure BP at 10, 14, 18, 21, and 22 weeks, according to the manufacturer’s recommended method. Measurements of diastolic and systolic pressure, mean pressure, heart rate, blood flow, and blood volume were obtained during echocardiogram sessions.(32) Measurements from de-identified rats were sorted at week 18 based on the ND or HFD diet of the rats, followed by RLX or placebo treatment.

### Optical Mapping

The optical mapping apparatus was previously described.(30) Briefly, hearts were excised and perfused on a Langendorff apparatus with Tyrode’s solution containing (in mM): NaCl (122), KCl (4.81), MgSO_4_ (2.75), NaHCO_3_ (25), Glucose 5), CaCl_2_ (2), gassed with 95% O_2_ and 5% CO_2_, pH 7.2 at 37°C. Hearts were placed in a custom-designed chamber to abate motion artifacts, and blebbistatin (5 µM) was briefly (15-20 min) added to the perfusate to minimize motion artifacts. Bolus injections of voltage (RH 237, 25 µl of 2 mg/mL dimethyl sulfoxide (DMSO) and Ca^2+^-indicator dye (Rhod-2/AM, 150 µl of 2 mg/mL DMSO) were made in the air-trap above the aortic cannula. Fluorescence images of Ca^2+^ transients (CaT) at 585±30 nm and action potential (AP) signals at >630 nm were recorded with 2 CMOS (Complementary Metal-Oxide-Semiconductor) cameras (Sci Media, Ultima One); each camera sensor (1cm x 1cm) recorded at 1000 frames/s with 100X100 pixels. The cameras were focused on the anterior surface of the ventricles or the left atrium (LA) to map APs and CaTs from the surface of the heart with a temporal and spatial resolution of 1ms and 250×250 µm^2^, respectively. The hearts were paced by 3 protocols: i) at various baseline cycle lengths (CL), ii) programmed electrical stimulation (PES): S1-S1=300 ms pacing at a baseline CL for 10 beats followed by a single S2 pulse delivered at decreasing S1-S2 intervals and iii) burst pacing (50 ms CL for 3s). Electrical stimuli were applied with a unipolar electrode on the edge of the right atrium and the base of the ventricles, respectively. Changes in conduction velocity (CV) without and with RLX treatment were measured at a baseline CL (S1-S1) or through the restitution kinetics of CV; that is, the CV of a single premature impulse delivered at decreasing S1-S2, until the premature impulse, S2 failed to capture at the refractory period or S2 initiated an arrhythmia.

### Antibodies

Validated antibodies were used to track changes in the expression of the following peptides: SCN5A guinea pig polyclonal antibody for Nav1.5 channels (Thermo Fisher Scientific, PA5-111796); Connexin 43 (Cx43) rabbit polyclonal antibody (Thermo Fisher Scientific, 71-0700); β-catenin rabbit monoclonal antibody (Abcam, ab32572); Wnt1 mouse monoclonal antibody (Thermo Fisher Scientific, MA5-15544); and Collagen I rabbit polyclonal antibody (Developmental Studies Hybridoma Bank from University of Iowa). The secondary antibodies were goat anti-Mouse Alexa Fluor 488 (Invitrogen, A-11029), a goat anti-Guinea Pig DyLight 488 (Invitrogen, SA5-10094), a goat anti-mouse Cy3 and goat anti-rabbit Cy3 secondary antibody (Jackson Immunoresearch, respectively: 115-165-003 and 115-165-144).

Immunofluorescence data was obtained from the base of the left ventricle (LV). Tissues were fixed with 2% paraformaldehyde made in PBS, equilibrated in 30% sucrose solution, and frozen in supercooled 2-methylbutane. LV cryosections (7µm thick) were labeled with Connexin 43, Nav1.5, β-catenin, Wnt1 or Collagen I antibodies by incubation with PBS-Triton-X-0.25% for 10 minutes, blocking with 2% BSA for one hour, and primary antibody incubation for 2 hours at 23 ^°^C. Appropriate secondary antibodies were applied for each primary antibody followed by DAPI staining. Imaging used an Olympus Fluoview 1000-2 and Olympus Fluoview 1000-3 Confocal microscopes (University of Pittsburgh, Cell Biology Imaging Center). Once optimal microscope settings were determined for each protein of interest, all subsequent images were taken with identical microscope settings. Two tissue sections per animal and three images per slide were captured. Post imaging analysis for relative expression was completed using FIJI ImageJ II software.

### Transcriptional Analysis

Heart RNA was isolated using Trizol Reagent (15596-026, Invitrogen). RNA was treated with DNAse I (E1010, Zymo Research) following the manufacturer’s instructions. RNA quality and concentration were determined on a BioDrop spectrophotometer (BD1800, Harvard Apparatus). Reverse transcription was performed using the MultiScribe Reverse Transcriptase System (43-112-35, Fisher). Sixteen ng of cDNA was used per qPCR reaction on a CFX Connect Real-Time System (CFX96, Bio-Rad) using PowerUP SYBR Green Master Mix (A25741, Applied Biosystems) as per manufacturer instructions. The amplification protocol was as follows: one cycle of sample denaturation at 95 °C (10 minutes), 40 cycles of denaturation at 95 °C (20 seconds), annealing at 58 °C (20 seconds), and elongation and acquisition at 72°C (1 minute). Relative expression was calculated using the 2^-ΔΔCt^ method with *RPLP0* as the housekeeping gene. Six biological samples were run in technical triplicate per condition. The rat Primers are listed in Table 1.

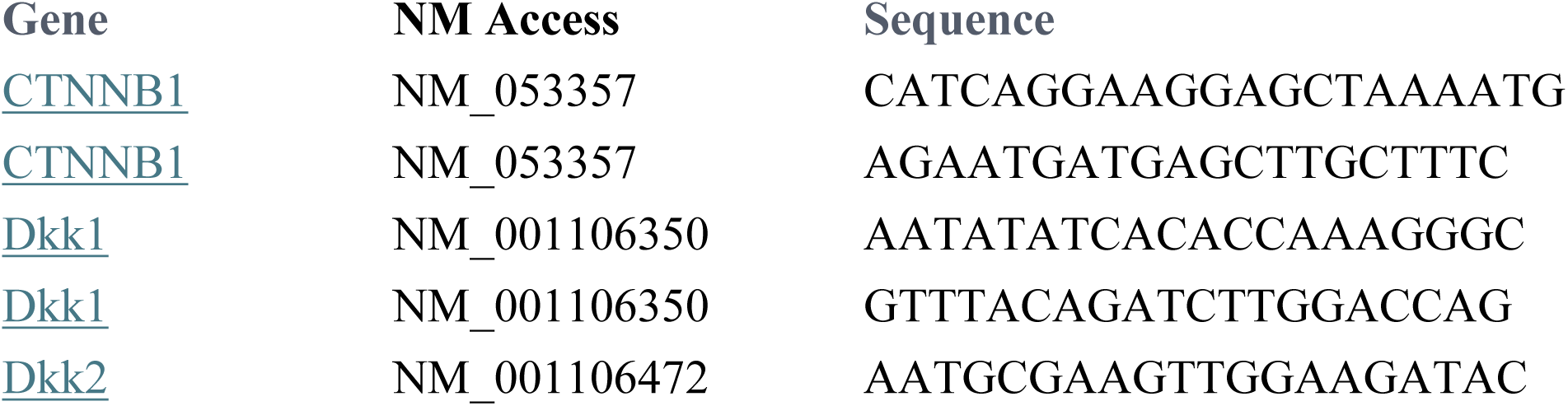
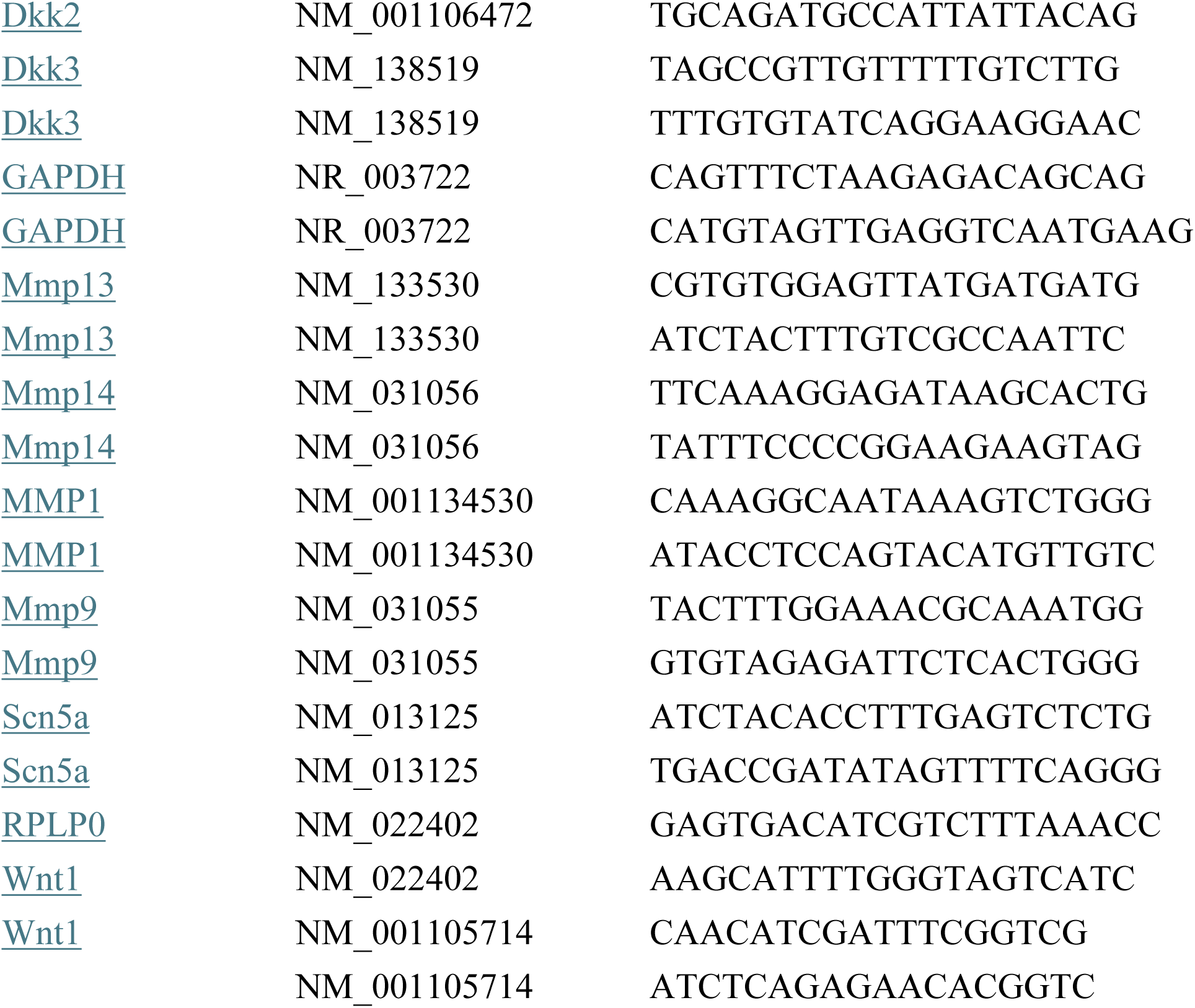

### Statistics

Statistical analyses were performed with GraphPad Prism 10.2.3 software (GraphPad Software). Expression analysis was performed using n=6 biological replicates and run in technical triplicates. Statistical comparisons between the two groups were performed using a nonparametric Mann-Whitney *U* test. The Shapiro-Wilk test was used for the normal Gaussian distribution of data. Data are presented as mean ± SD.

### Serum Biomarker Quantification

Blood obtained before the optical mapping was immediately placed in non-heparinized collection tubes for serum collection. Serum samples from each rat were sent to Relaxera in Bensheim, Germany for quantification of circulating levels of RLX, and NT-pro-ANP. All kits were used according to the manufacturers’ instructions (RLX, Quantikine ELISA kit [R&D Systems, DRL200]; rat NT-pro-ANP [BIOMEDICA, Vienna, Austria, catalogue no. 20892]

## Results

### Determination of HFpEF in ZSF1 Rats

The pathology of HFpEF is manifested at multiple organs, and here, we based the HFpEF diagnosis through ultrasound measurements and a biomarker of hemodynamic stress (NT-pro-ANP) in intact animals followed by invasive tests of arrhythmia vulnerability, changes in gene and protein expression.(33) Renal function was not analyzed, as it had been previously investigated in the same rat model for HFpEF.(14) Figure 1 shows the change of various echocardiogram parameters from week 10 to week 18 in ZSF1-obese rats receiving a HFD. Ejection fraction (EF) is maintained well above 50% up to week 18, indicating a preserved systolic function (Figure 1B). Early diastolic mitral annular velocity, e’, is measured to reflect LV diastolic function.(34) At week 18, e’ was about -36 mm/s, a significant decrease in absolute magnitude from week 10, at a baseline value of -50 mm/s (Figure 1C). The ratio of early mitral inflow velocity and early diastolic mitral annular velocity (E/e’), a measure of LV filling pressure and conveys diastolic function.(35) By week 18, the animals on HFD displayed an E/e’ of about -25; a significant increase in absolute magnitude from the week 10 baseline value of -18 (Figure 1D). Cardiac hypertrophy was also apparent as the left atrial length measured using a parasternal long axis view (PSLAX), increased from about 4.1 mm at baseline to 5.1 mm by week 18 (Figure 1F), and LV posterior wall diameter (LVPW d) increased significantly from 20.3 mm at baseline to 25.8 mm by week 18 (Figure 1G). LV mass significantly increased from a baseline of ∼915 mg to ∼1203 mg by week 18 (Figure 1H). LV mass was corrected for overestimation according to previously established guidelines.(36) Lastly, cardiac index (CI) was used as a measure of cardiac output, as weight is an important factor in HFpEF development, CI decreased significantly from 212.2 at week 10 to 156.2 at week 18 (Figure 1I).

**Figure 1:**
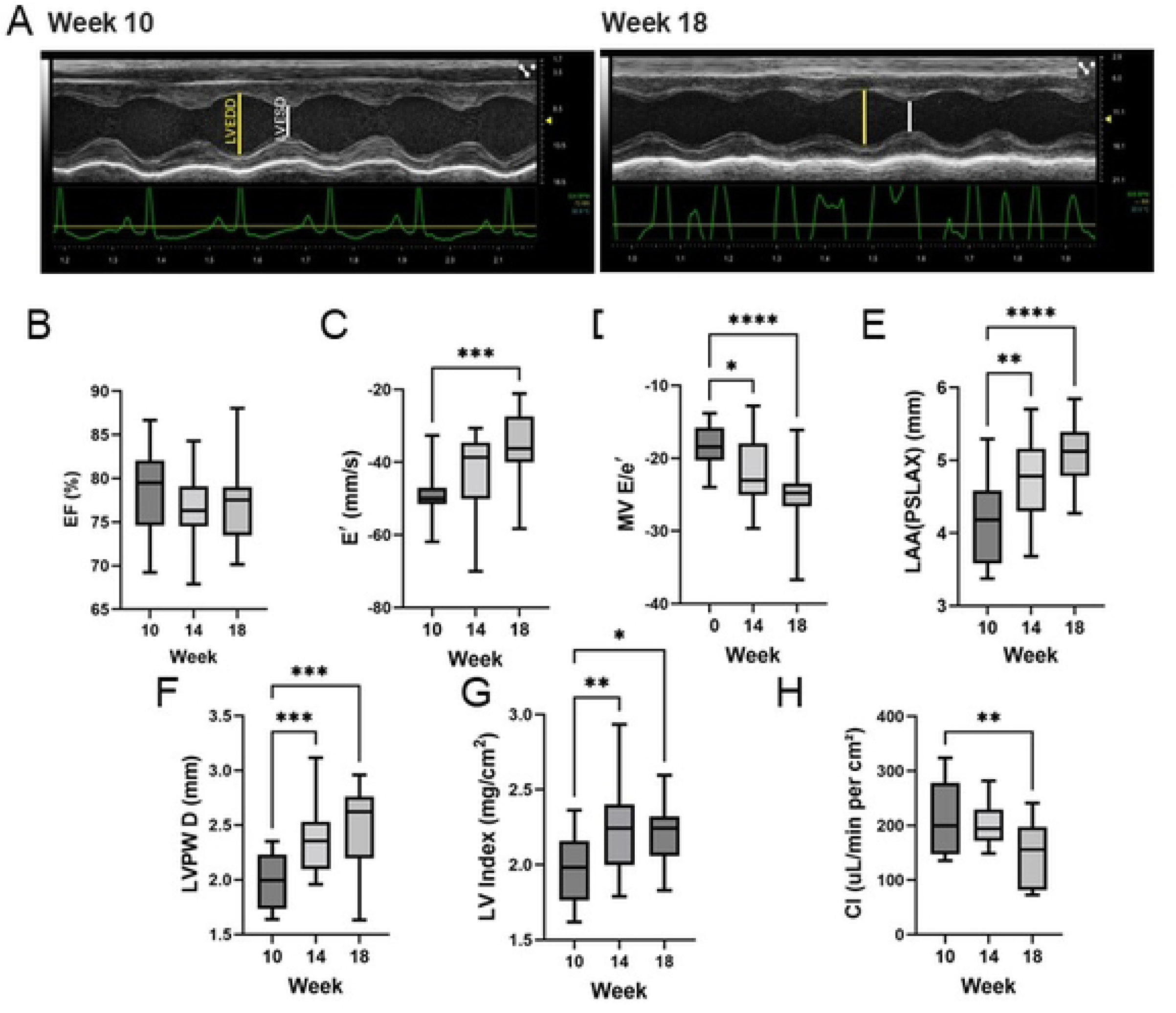
Ultrasound parameters used to track the development of HFpEF, week 10 to 18. (A) Ultrasound images used to identify changes in left ventricular end diastolic diameter (LVEDD) and left ventricular end systolic diameter (LVESD), at week 10 (left) and week 18 (right). Quantitative comparison of key echocardiogram parameters for weeks 10, 14 to 18, during HFpEF development. (B) Percent ejection fraction (EF). (C) Early diastolic mitral annular velocity (e’). (D) Ratio of early mitral inflow velocity and early diastolic mitral annular velocity (E/e’). (E) Parasternal long axis view of left atrium size (PSLAX). (F) Left ventricular posterior wall diameter. (G) Left ventricular mass index. (H) Cardiac Index (CI). ZSF1-obese rats were placed on a HFD (High Fat Diet, N=11) or a ND (Normal diet, N=8) from week 10-18, then were implanted with osmotic mini-pumps at week 20 and continued on the same diet till week 22. The mini-pumps delivered either the VEH (Controls) or RLX; in all panels, error bars indicate minima and maxima from the mean. Grubbs outlier test was used to remove statistical outliers (p<0.05). ** indicates p<0.01, *** indicates p<0.001, **** indicates p<0.0001.

### Effects of Relaxin on Diastolic Dysfunction in HFpEF

Echocardiogram measurements after two weeks of RLX or VEH administration were compared to those from week 18, when the HFpEF diagnosis was established. EF remained above the 50% threshold for HFpEF in RLX and VEH groups, still EF experienced a significant drop from week 18 to week 22 in VEH treated rats but the drop was not significant in RLX-treated rats (Figure 2B). Interestingly, the key measures of diastolic function, e’, MV E/e’, and LAA were significantly improved by RLX-treatment (Figure 2 C, D, F). This suggests that RLX-treatment improved diastolic function in this model of HFpEF.(3) The remaining parameters (Figure 2 F-H) did not change in either RLX or VEH treatment.

**Figure 2:**
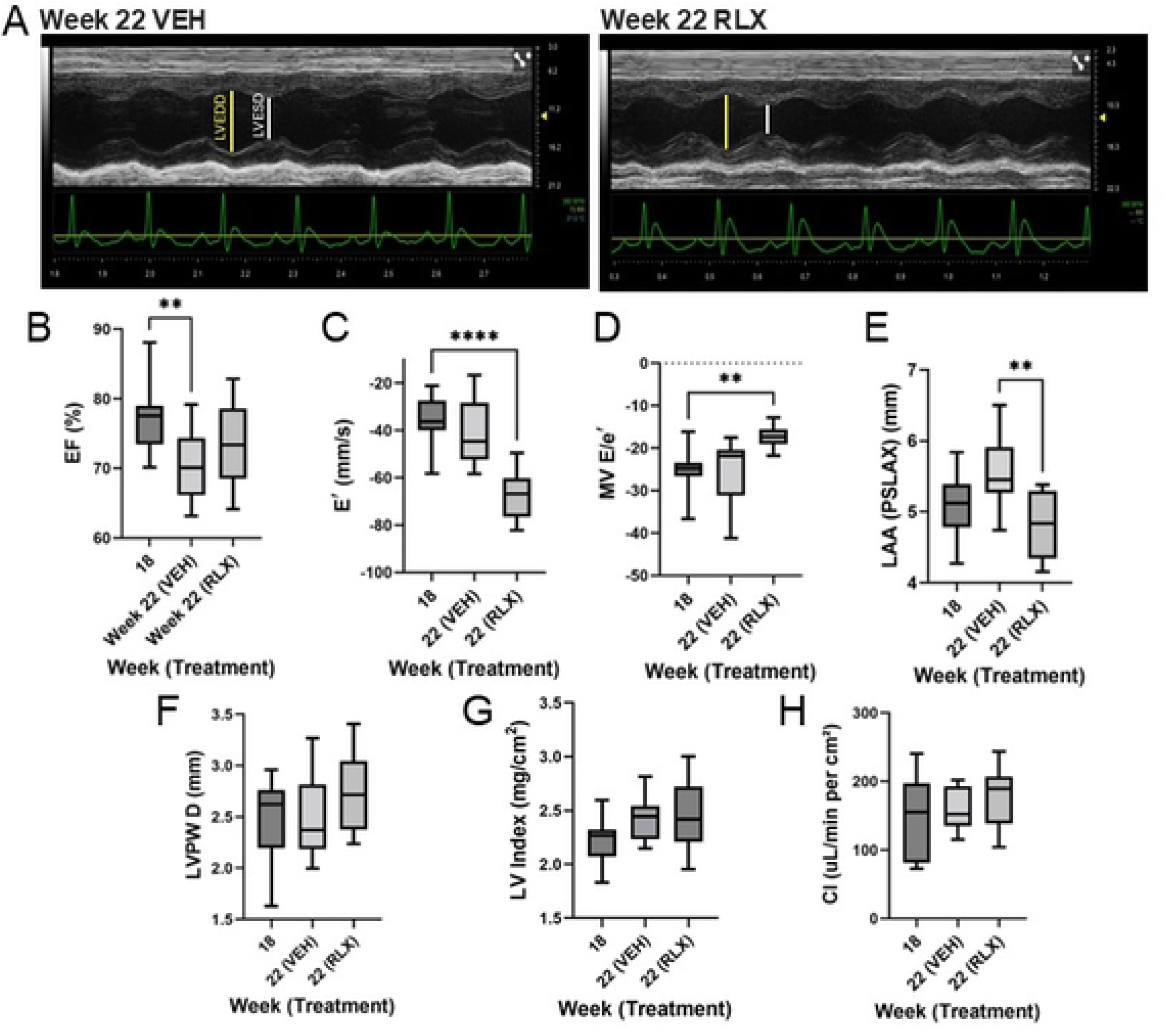
Effects of Relaxin Treatment on HFpEF by Echocardiogram Analysis. (A) Ultrasound images identifying left ventricular end diastolic diameter (LVEDD) and left ventricular end systolic diameter (LVESD) in HFpEF rats treated with the Vehicle (VEH) or Relaxin (RLX) (right). Comparison of echocardiogram parameters in HFpEF rats at week 18, followed by treatment with VEH or RLX from week 20 to 22. (B) Percent ejection fraction. (C) Early diastolic mitral annular velocity (e’). (D) Ratio of early mitral inflow velocity and early diastolic mitral annular velocity (E/e’). (E) Parasternal long axis views of left atrium size (PSLAX). (F) Left ventricular posterior wall diameter. (G) Left ventricular mass index. (H) Cardiac Index. Animals were either on a HFD (n=11) or on a ND (n=8) from week 10 to week 22, error bars indicate minimum and maximum in all panels. The Grubbs outlier test was used to remove statistical outliers (p<0.05). ** indicates p<0.01, **** indicates p<0.0001.

The circulating levels of NT-pro-ANP are a measure of hemodynamic stress that occur in HFpEF and after RLX therapy. In healthy control rats, the median NT-pro-ANP was 0.3 nmol/L (N=10, with an interquartile range (IQR)= 0.22-0.55) and increased to 1.59 nmol/L (N=3, IQR=1.23-2.06) in ZSF1 rats placed on a HFD. RLX-treatment suppressed the elevation of NT-pro-ANP to 0.5 nmol/L (N=4, IQR=0.43-1.03); this lowering of NT-pro-ANP was not seen in HFpEF rats that received the placebo. Consistent with a previous report,(37) ZSF1-obese rats increased their body weight by 38%, from 420 to 580 grams in 11 weeks, irrespective of treatment (HFD vs ND ± RLX-treatment and exhibited mild hypertension with diastolic and systolic pressures in the range of 100-120 mmHg and 140-160 mmHg, respectively.

### Effects of Relaxin on Atrial and Ventricular Arrhythmia Phenotype

ZSF1-obese rats placed on a normal diet (ND) did not develop HFpEF and served as controls for rats placed on a high-fat diet (HFD) that developed HFpEF. The susceptibility to AF and VF was tested by PES. Figure 3A illustrates an example of a control heart, first paced at a cycle length of 300 ms, then 80 ms for 2 sec resulting in a brief atrial arrhythmia lasting 2 sec that self-terminated back to sinus rhythm. In contrast, control hearts treated with RLX did not exhibit atrial arrhythmias, either transient or sustained (Fig. 3B). In contrast, HFpEF rats were prone to sustained atrial arrhythmias (Fig. 3A’) in 30% of the hearts, after a single premature impulse at S1-S2=20 ms. HFpEF rats treated with RLX did not exhibit sustained AF even after burst pacing on the atria (Fig. 3B’). Atria of RLX treated animals tended to have slightly higher conduction velocities than VEH treated (Fig, 3D’). The arrhythmia profile (Fig. 3C’) shows that RLX treatment blocked sustained AF and allowed transient self-terminating atrial arrhythmias in 1/3 of the hearts, whereas 1/3 of VEH treated rats had sustained AF, self-terminating or no arrhythmias (Figure 3C’). In untreated rats, the propensity to no arrhythmia, sustained and un-sustained arrhythmias were equal but after RLX-treatment, sustained atrial arrhythmias were not detected and only brief arrhythmias lasting <10 s were observed.

**Figure 3.**
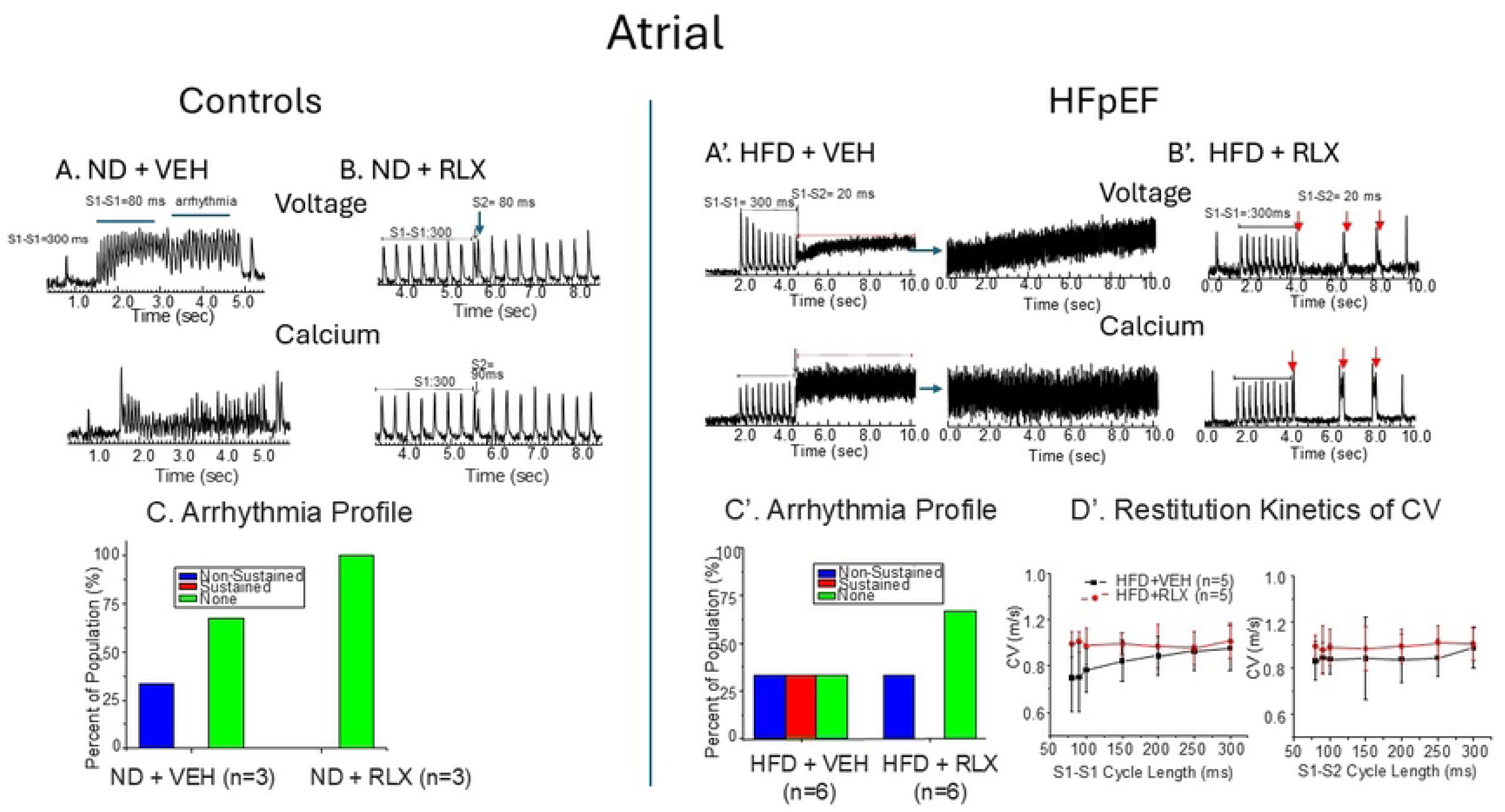
Atrial Arrhythmia vulnerability in Control and HFpEF rats ± RLX treatment. Optical action potentials (APs) and Ca^2+^ transients (CaTs) were simultaneously recorded from the Right Atrium (RA). Panel A: An example of atrial APs (top) and CaTs (bottom) that are recorded simultaneously from a site on the RA of a control, non-HFpEF rat heart placed on a normal diet (ND) and treated with VEH (control). Rapid pacing at S1-S1= 80 ms for 2 sec, resulted in a transient, non-sustained, self-terminating tachycardia for ∼ 2 sec. Panel B: Rats on a ND were treated with RLX and a premature impulse at S1-S2 = 80 ms resulted in a single extra pulse and a return to normal sinus rhythm. Panel C: Summary of the propensity to arrhythmia: No arrhythmias (blue), sustained AF (red) and non-sustained arrhythmias, lasting <10 sec (Green). The profile showed that 1/3 of non-HFpEF rats had transient atrial arrhythmia and 2/3 had no arrhythmia. RLX suppresses transient arrhythmia. Panel A’: A premature impulse at S1-S2 = 20 ms elicited a sustained AF in HFpEF rats which lasted for the duration of the experiment. Panel B’: The same pacing protocols applied to a HFpEF rat treated with RLX failed to elicit an arrhythmia and repeated premature impulse either elicited small extra APs (red arrows) or failed to capture. Panel C’: As in panel C for HFpEF hearts. The arrhythmia profile showed that in VEH treated HFpEF rats, a premature impulse elicited either no atrial arrhythmias (1/3), or non-sustained (1/3) or sustained arrhythmia (1/3). In RLX treated HFpEF rats, a premature impulse elicited a brief arrhythmia lasting < 10 sec or no arrhythmia. Panel D’: In RLX treated hearts, atrial CV tended to be faster than in VEH treated atria, an effect found to be more pronounced at faster heart rates or shorter cycle lengths in the range of 100-150 ms. CVs were measured while pacing at cycle lengths 100 to 300 ms (left graph) or the CV of a single premature impulse elicited by S1-S2 from 75-250 ms.

Similar trends were observed with ventricular arrhythmias (Fig. 4). ZSF1-obese rats on a ND and no HFpEF, all had brief, self-terminating ventricular fibrillation (VF) (Fig. 4A), and RLX-treatment reduced the incidence of transient arrhythmias to 25% (Fig. 4B) and the rest had no arrhythmia. In HFpEF hearts from VEH treated animals, 1/3 experienced sustained VF (Fig. 4A’), 1/3 non-sustained VF, and 1/3 no arrhythmia (Fig. 4C’). RLX-treatment blocked sustained VF (Fig. 4B’) and the incidence of transient, self-terminating arrhythmias was at 50% of the hearts (Fig. 4C’). In HFpEF hearts, RLX caused a marked increase in the CV of premature impulses (Fig. 4D’) which can account for its anti-arrhythmic actions. These data indicate that RLX-treatment of rats with HFpEF is protective of atrial and ventricular arrhythmia and effectively blocks sustained AF and VF.

**Figure 4.**
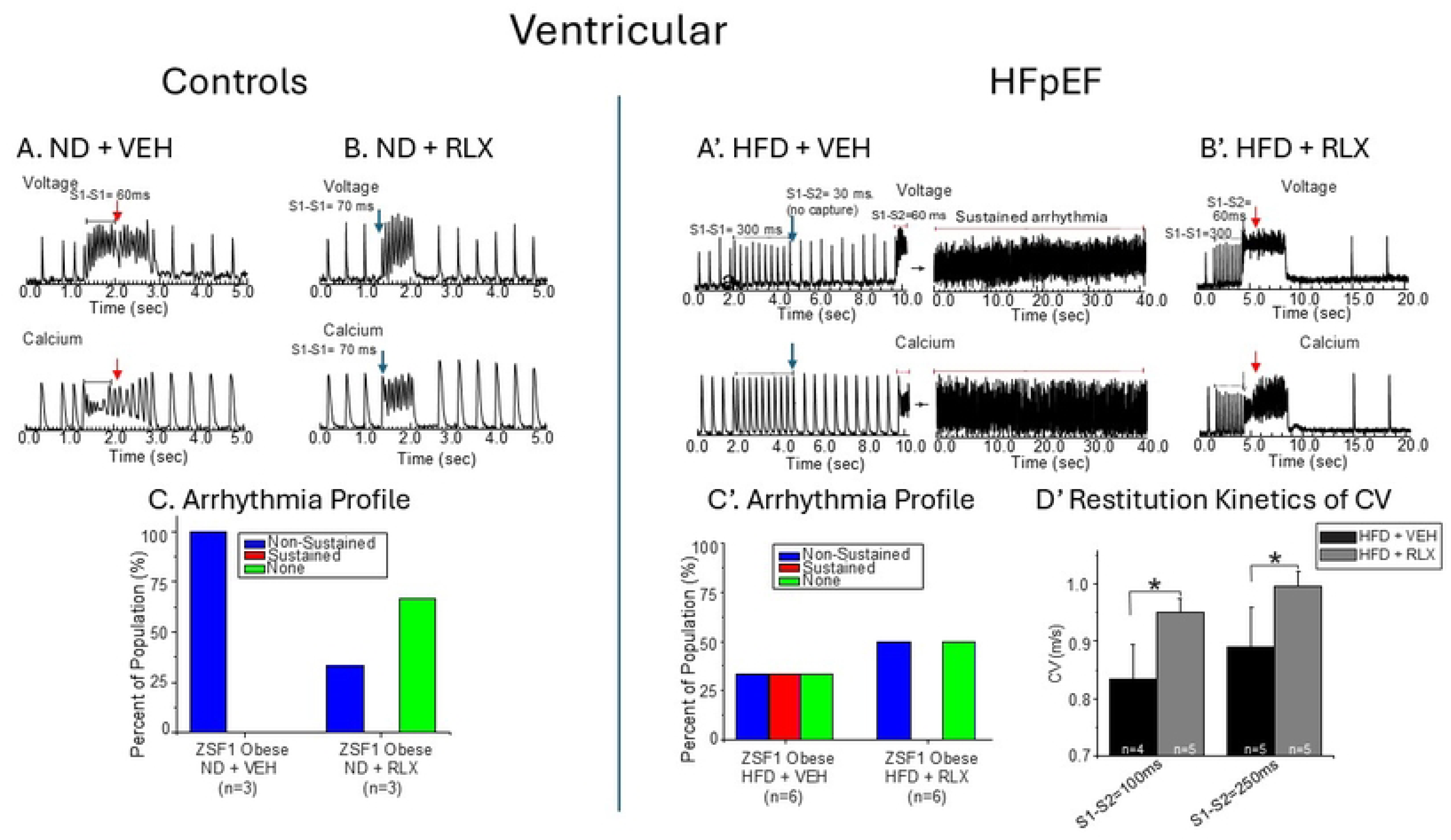
Ventricular Arrhythmia vulnerability in Control and HFpEF rats ± RLX treatment. Panel A: ZSF1-obese rats on a ND had brief, non-sustained arrhythmia following rapid pacing. Panel B: RLX-treatment reduced the vulnerability to non-sustained ventricular arrhythmia. Panel C: Summary of the propensity to arrhythmia: No arrhythmias (blue), sustained AF (red) and non-sustained arrhythmias, lasting <10 sec (Green). The ventricular arrhythmia profile showed that 3 ZSF1-obese rats exhibited non-sustained ventricular arrhythmia following rapid pacing or a premature impulse. In RLX treated hearts, arrhythmias could not be elicited by rapid pacing or premature stimuli, in 2/3 of the hearts. Panel A’: Illustrates a train of APs and CaTs during pacing at a basic CL of 300 ms, followed by a premature impulse at 30 ms (S1-S2 = 30 ms) which failed to capture and a premature impulse at 60 ms (S1-S2 = 60 ms) which elicited a sustained VF, lasting for the duration of the experiment. Panel B’: Illustrates a HFpEF hearts treated with RLX, first paced at a CL = 300 ms and a premature impulse at 60 ms elicited a non-sustained, self-terminating VF that lasted ∼ 3 sec. Panel C’: As in panel C. The ventricular arrhythmia profile shows that HFpEF hearts exhibited equally 2/6 no arrhythmia, non-sustained and sustained arrhythmia. RLX treated HFpEF hearts exhibited transient arrhythmia (3/6) or no arrhythmia (3/6). D’: The CV of premature impulses were measured in HFpEF hearts with VEH or RLX treatment. RLX treatment resulted in a statistically significant increase in CV at S1-S2 = 100 ms and 250 ms, which is consistent with a reduced susceptibility to reentrant arrhythmia.

### Effects of RLX on myocyte remodeling

By analogy with our previous studies on aged rats,(19, 22, 29) RLX-treated HFpEF rats had a significantly greater expression of the gap-junction protein Connexin 43 (23.3%) (Fig. 5A) and Nav1.5 (29%) (Fig. 5B) compared to VEH treated rats. In addition, Cx43 tended to be more localized to the lateral membrane in HFpEF rats compared to controls, and RLX increased the localization of Cx43 to intercalated disks (Fig. 5A). However, there was no significant difference in Cx43 in ZSF1 rats on normal diets treated with VEH or RLX. Interestingly, RLX treatment of control rats on a normal diet expressed higher levels of Nav1.5 compared to VEH treated control rats and RLX treated HFpEF rats. These changes in gene expression induced by RLX impart the heart with improved cell-cell coupling and conduction velocity which reduces the vulnerability to arrhythmias.

**Figure 5.**
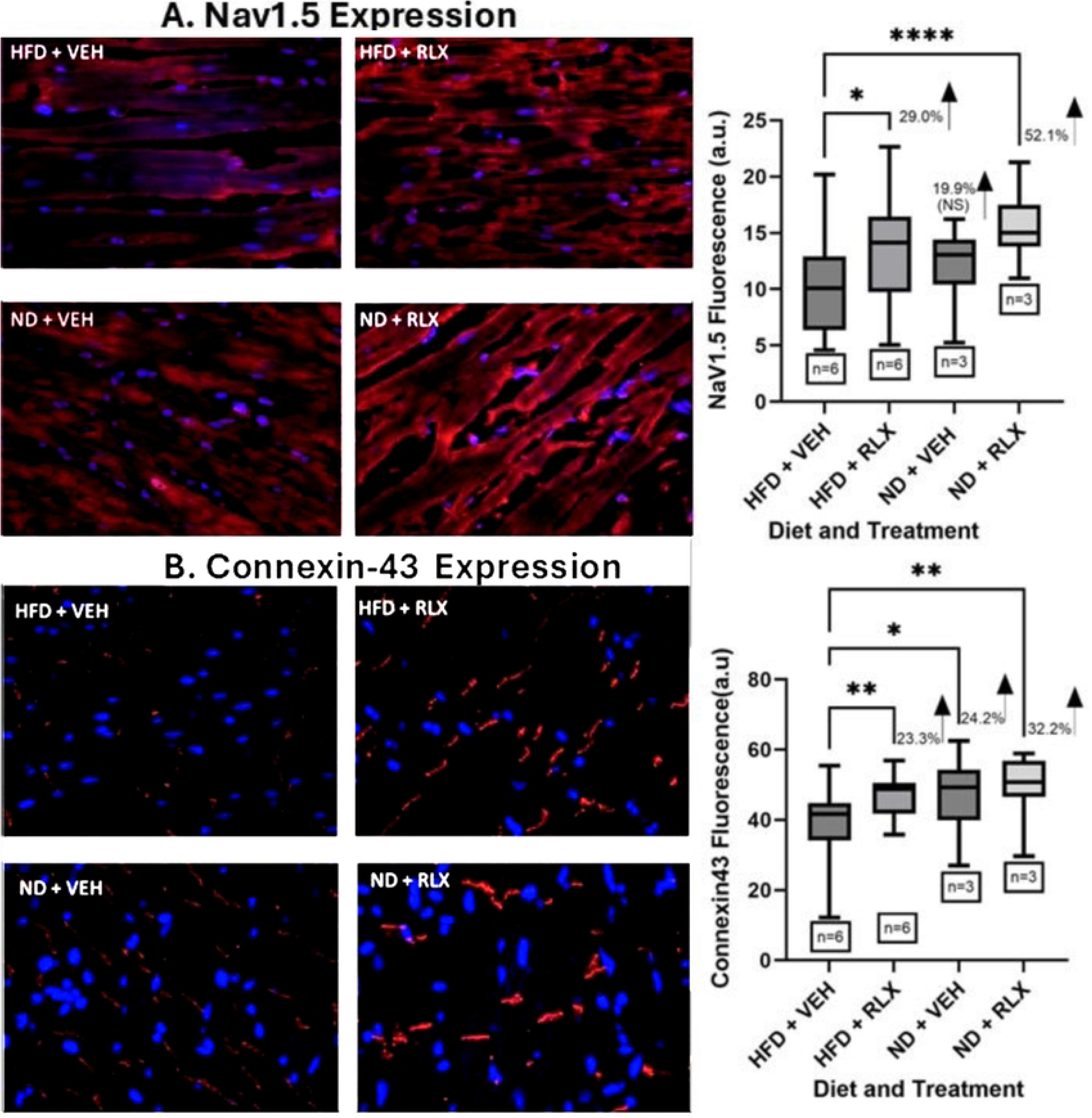
Ion channel remodeling in Hearts after RLX Treatment. Hearts from the rats studied in Figures 1-4 were frozen and tissue sections were used to analyze changes in protein expression by immuno-fluorescence (IF) in Figures 5-7. (A) RLX markedly increased Nav1.5 expression in HFpEF hearts compared to VEH treated controls (top right vs. top left panels). A right, bar graphs: Quantification of Nav1.5 expression in hearts on a HFD and a ND with VEH or RLX treatment. (B) Cx43 expression was the lowest in HFpEF control hearts (top left) and RLX treatment increased Cx43 (top right) to levels in hearts on a ND (bottom left). RLX also shifted the localization of Cx43 from the lateral membrane to intercalated disks; compare (top left to top right). B, bar graphs: Quantification of Cx43 expression in hearts on a HFD and a ND with VEH or RLX treatment. Error bars indicate SEM, N=6 rats per group with 6 images taken per rat for all VEH and RLX data and apply to all panels. ** indicates p<0.01. Images taken on Olympus Fluoview Confocal Microscope at 60x magnification.

Based on the role of Wnt1 canonical signaling in the actions of RLX in aged hearts, we investigated the expression of β-catenin and its localization at intercalated disks of myocytes.(19, 38) RLX increased Wnt1 expression on average by 36.2% in rats with HFpEF but had no significant effect on control hearts ND (Fig. 6A). Consistent with the activation of Wnt canonical signaling, RLX increased β-catenin expression (33.0%) in HFpEF hearts with no significant effect in control hearts (Fig. 6B). The antifibrotic actions of RLX demonstrated in previous investigations were measured by measuring collagen I levels and picrosirius red labeling.(22) In this HFpEF model, fibrosis measured with a collagen I antibody (Fig. 7 A) or picrosirius red (Fig. 7B) was not significantly altered in ZSF1 hearts placed in a ND or a HFD with or without RLX treatment.

**Figure 6.**
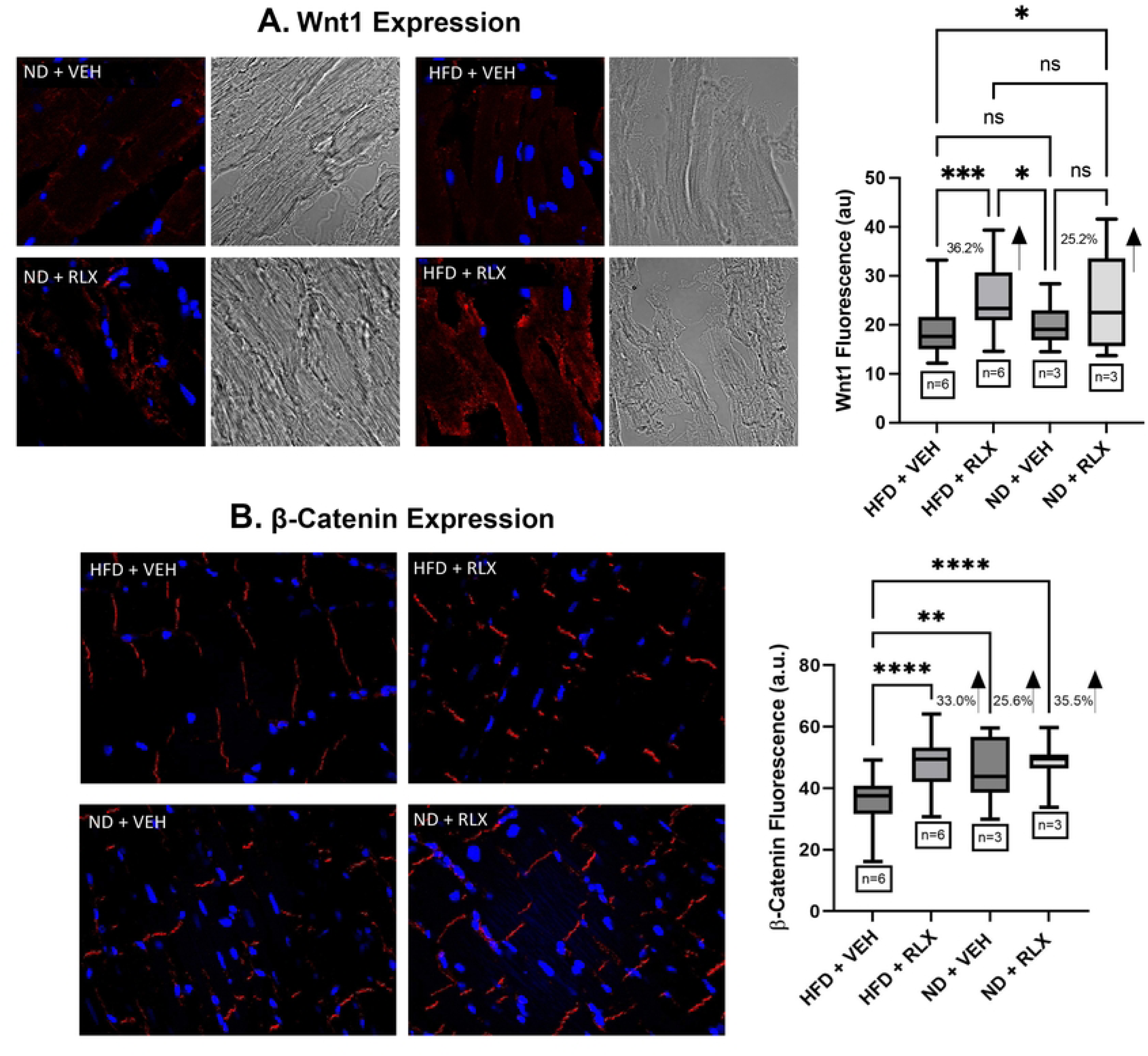
RLX treatment activates the Wnt/canonical signaling pathway consistent with genomic modifications of HFpEF rats. (A) IF quantification of Wnt1 expression as a function of diet (ND or HFD; that is, non-HFpEF or HFpEF rats) and treatment (VEH or RLX). Top panels: Confocal images of ventricular tissue sections from non-HFpEF rats labeled with DAPI and a Wnt1 antibody and adjacent transmitted light images of the same section to better visualize individual cells. RLX treatment did not significantly alter Wnt1 expression. Bottom panels: Confocal sections as in top panels from HFD treated rats that develop HFpEF; here, RLX increased Wnt1 expression in intracellular vesicles within myocytes. A right, bar graphs: Statistical analysis of Wnt1 expression in hearts from rats placed in HFD or ND then treated with VEH or RLX. (B) IF quantification of β-catenin expression as a function of diet and treatment post-HFpEF. (B right, bar graphs: Statistical analysis of β-catenin expression in hearts from rats placed in HFD or ND then treated with VEH or RLX. RLX treatment markedly increased β-catenin and Wnt1 in HFpEF rats, indicative of the activation of Wnt canonical pathway. All data across groups obtained from slides stained under same conditions. Error bars indicate SEM. All data from ND animals unless specified with HFD, N=3 rats per group for normal diet and 6 rats per group for HFD with 6 images analyzed per rat which applies to all panels. * Indicates p<0.05, ** indicates p<0.01. Images taken on Olympus Fluoview Confocal Microscope at 60x magnification.

**Figure 7.**
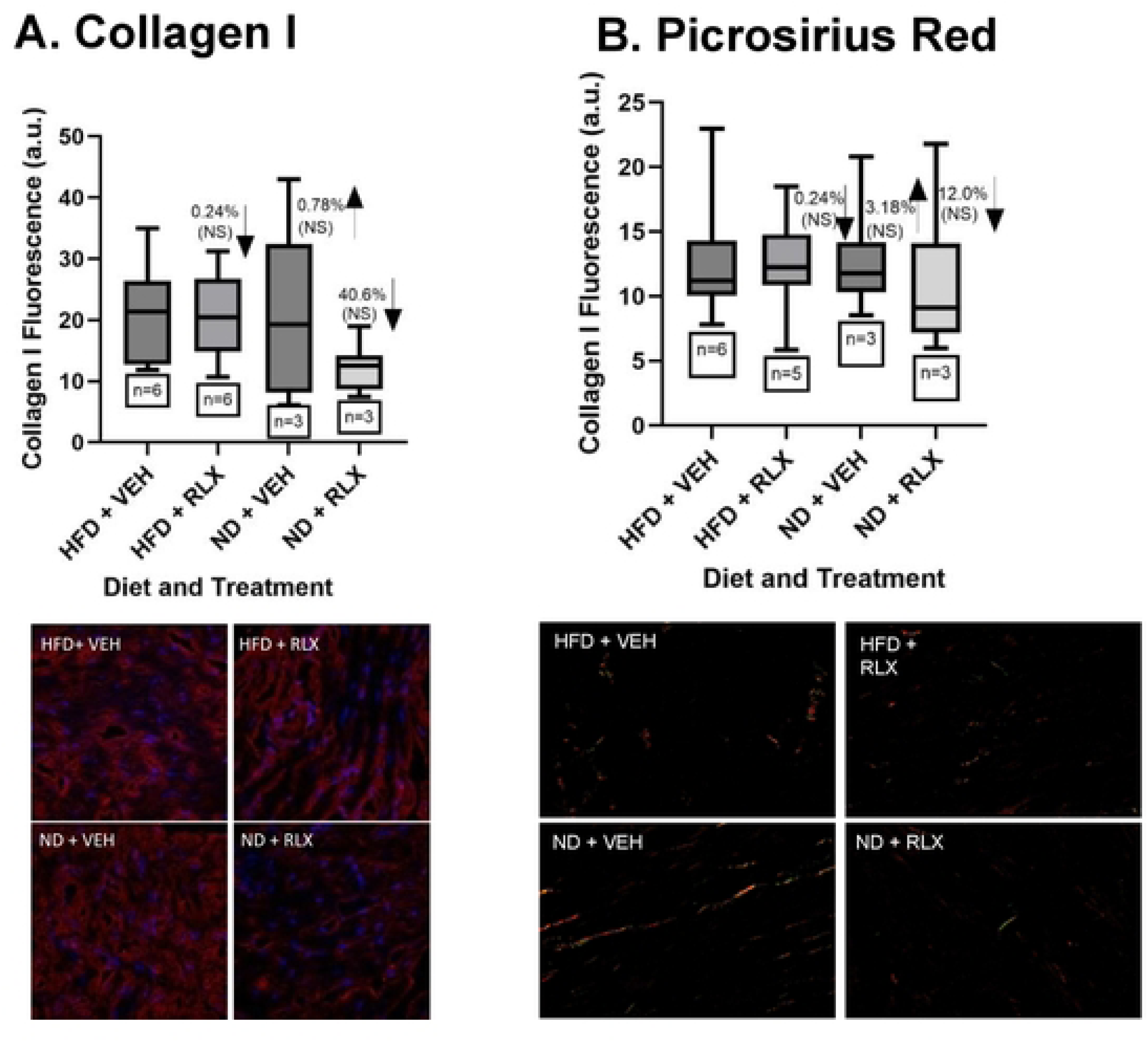
Propensity to Fibrosis in ZSF1-obese rats. The role of fibrosis in the ZSF1-obese rat model of HFpEF was examined using an antibody against collagen 1 (A) and picrosirius red (B). There were no significant changes in fibrosis between ZSF1-obese rats on a ND or a HFD and RLX treatment did not alter the density of fibrosis in the ventricles.

### qPCR Analysis

Based on our previous studies on the genomic effects of RLX on aged rat hearts,(22, 28) selected genes were investigated by qPCR. RLX was shown to activate Wnt canonical pathway (Wnt1 and CTNNB1), suppress Dickkopf peptides (endogenous inhibitors of Wnt signaling: DKK1, 2, and 3), and alter metalloproteases predominantly found in the heart (MMP1A, MMP9, MMP13 and MMP14). Figure 8 summarizes some changes in gene expression caused by HFpEF with and without RLX-treatment. Significant decreases in gene expression were found in DKK1, and MMP1A in RLX compared to the VEH treated HFpEF group, while SCN5 shows a significant increase in expression.

**Figure 8:**
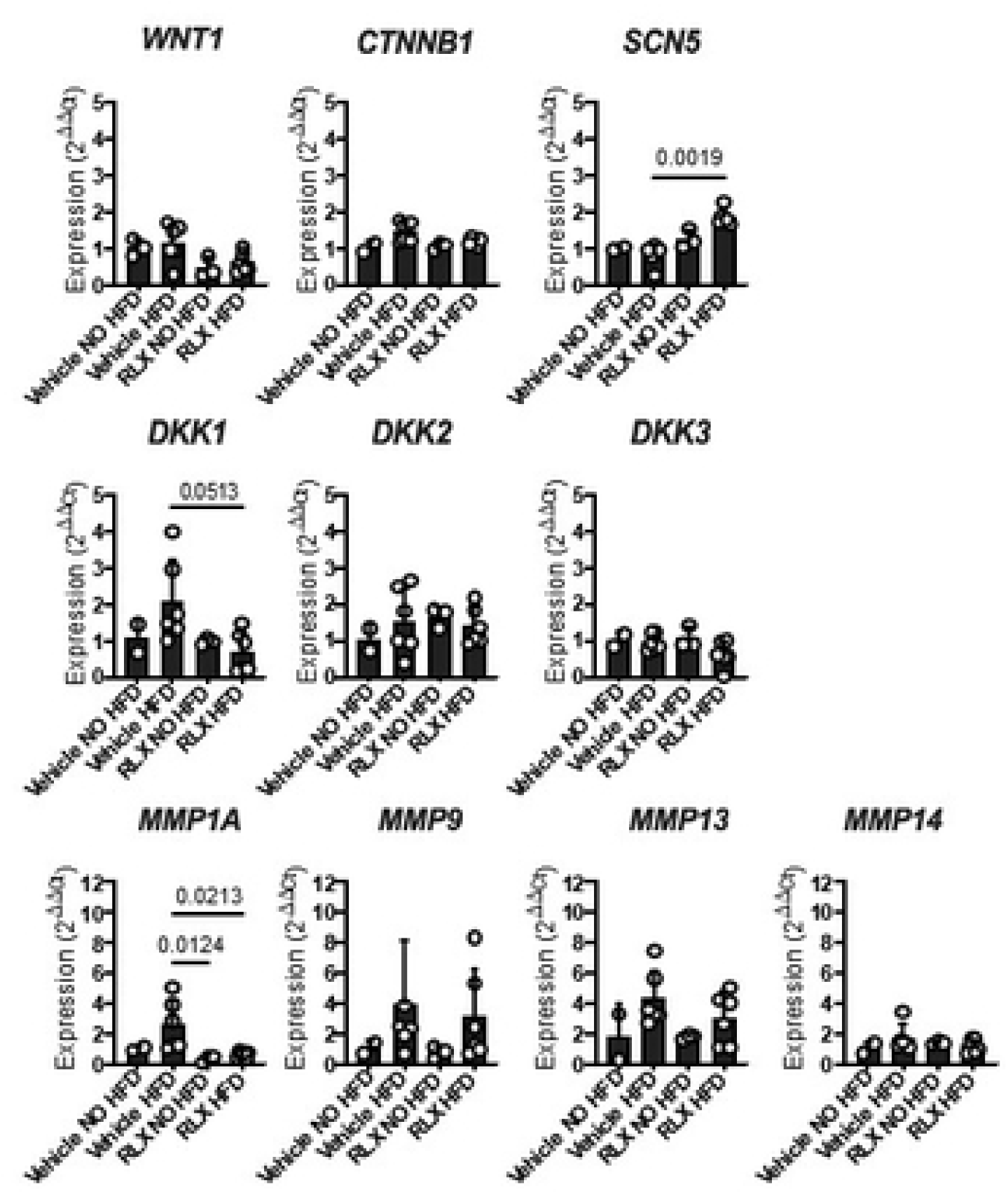
Expression profile changes in HFpEF rats with and without RLX treatment. Male ZSF1 rats were randomly assigned to either a control group or a high-fat diet (HFD) for 20 weeks, followed by 2 weeks of RLX or vehicle administration. After collecting blood samples, recording weight, and conducting echocardiographic measurements, the hearts were isolated, perfused using a Langendorff apparatus, and frozen for RNA extraction and qRT-PCR analysis. Expression is shown as relative expression, which was calculated using the average threshold cycle number and shown as the 2 ^(Ct(*housekeeping* gene)−Ct(*target gene*))^. Gene names are displayed at the top of each graph. n = 3 in the non-HFD groups , n = 6 in the HFD groups. Data are presented as means ± SD. *P-*values were calculated using the Kruskal-Wallis H test and Dunn’s pairwise comparison post hoc tests, with significant values indicated on the graphs.

## Discussion

The ZSF1-obese rat on a HFD developed many hallmarks of clinical HFpEF: obesity, hyperglycemia, DD, enlarged LA, high blood pressure, enhanced vulnerability to arrhythmia and hemodynamic stress (higher NT-pro-ANP). As previously described by Hamdani et al., these changes occurred in ZSF1-obese rats on a HFD gradually over a period of several weeks, while preserving a near-normal ejection fraction (EF>50%).(14) Here, we show that the continuous delivery of RLX via an osmotic minipump for 2-weeks greatly improved the diastolic compliance and reversed the LA enlargement and hemodynamic stress by decreasing circulating NT-pro-ANP, in intact animals. In isolated Langendorff perfused hearts, the prior RLX-treatment suppressed the vulnerability to sustained AF and VF triggered by PES, and the cardiac arrhythmia protection was primarily mediated by an increase in conduction velocity through increases in Nav1.5 and Cx43 expression. RLX is known to act via its cognate GPCR, RXFP1 and to exert its beneficial and therapeutic effects in the cardiovascular system by causing elevations in cAMP and cGMP.(39) The pleiotropic actions of RLX were initially studied through RXFP1-dependent mechanisms and did not address the possible cellular genomic modifications elicited by RLX. The expectation of genomic modifications was compelling given that in the RELAX-AHF-2 clinical trial, a 48-hour infusion of RLX (i.v.) reduced overall cardiovascular death by 37% 180 days post-treatment and that RLX is short-lived (∼2.5 hrs.).(40)

In aged hearts, we showed that RLX suppressed arrhythmias by increasing Nav1.5 and Connexin 43 expression by activating canonical Wnt signaling (increased Wnt1 and nuclear β-catenin), a mechanism that was blocked by exogenous Dickkopf-1(DKK1).(19) This fundamental mechanism is now shown to apply in HFpEF hearts, as RLX treatment increased Nav1.5 and Cx43, cytosolic Wnt1and β-catenin. RLX treatment also decreased mRNA levels of DKK1 with no effect on DKK2 or DKK3, which is an important component of Wnt/canonical signaling and congruent with previous findings on aged rats.(19)

These findings align with previous studies and confirm the genomic remodeling capabilities of RLX in heart diseases.(29) Consistent with previous reports,(14, 37) we found that this model of HFpEF doesn’t develop cardiac fibrosis and in the absence of excess fibrosis in HFpEF hearts, RLX-treatment had no effect on collagen deposition.

RLX treatment had no significant effect on blood pressure (BP), which is consistent with reports on spontaneously hypertensive rats and Angiotensin II-induced hypertensive rats, despite increases in systemic arterial compliance.(25, 41) In contrast, other studies reported that RLX treatment had a tendency to reduce BP(24), or that RLX caused a statistically significant decrease in BP in spontaneously hypertensive rat(42) and in angiotensin-II induced model of hypertension.(43) These contradictory findings require further investigation. We would like to emphasize, however, that all observed RLX effects are unrelated to BP, which is highly desirable for the clinical setting. Eventually, the neutral BP effect of RLX may account for the unchanged cardiac hypertrophy or Cardiac index after treatment.

In this study, RLX is not used as a prophylactic; the RLX treatment was applied to the animals after the development of hallmarks of HFpEF and a treatment of merely 2 weeks reversed the diastolic dysfunction (even as the rats were kept on the HFD), the LA enlargement, and the hemodynamic status. Another major action of RLX is the suppression of sustained arrhythmias, including AF triggered by a premature impulse or burst pacing. The latter is significant given that AF is particularly common in HFpEF patients with a prevalence of 40–60% and its occurrence heralds worsening of prognosis.(44) Furthermore, both in the EMPEROR-Preserved and DELIVER phase III trials investigating SGLT2 inhibitor therapy for HFpEF, sudden cardiac death remained the most important single cause of cardiovascular mortality.(44)

The qPCR analysis confirms the increase in Nav1.5 found in other studies, and the decrease in Dkk1 gene expression will also lead to an increase in Wnt signaling. The reduction in β-catenin is somewhat expected because β-catenin mRNA is primarily regulated by post-translational control.

## Limitations

The study did not investigate ZSF1-obese female rats on a HFD and the actions of RLX-treatment, although a study found that females also develop HFpEF with essentially similar pathology between the sexes.(37) The RLX treatment (dose and duration) was determined from previous studies for different conditions and additional doses and durations of treatment should be tested, specifically for HFpEF. The protocol included the analysis of the arrhythmia phenotype in Langendorff perfused hearts by optical mapping and of changes in protein expression in ventricular tissues by IF; follow up studies are needed to determine how long the RLX treatment imparts its beneficial effects, post treatment, the efficacy of prolonged RLX treatment and can smaller doses of RLX sustain the beneficial effects? Future studies will include both sexes and additional biomarkers of interest, such as circulating levels of NT-pro-BNP. Nevertheless, the study provides compelling evidence in support for RLX-treatment as a therapy for HFpEF.

## Conclusions

ZSF1 obese rats on HFD developed HFpEF, as verified by echocardiography, NT-pro-ANP, glucose, and systolic blood pressure elevation, in intact animals and by an enhanced vulnerability to arrhythmias in Langendorff perfused hearts. Human synthetic RLX infusion for 2 weeks via osmotic minipumps improved diastolic dysfunction. RLX also reversed maladaptive cardiac remodeling in HFpEF via reduced Dkk1 and increased β-catenin expression, implying an increase in Wnt/canonical signaling. This is consistent with the actions of RLX in aged rats which included an increase in the expression of Wnt1, β-catenin, Cx43, Nav1.5 and a decrease in collagen deposition.

## Acknowledgements

This publication was supported by the Samuel and Emma Winters Foundation, Department of Medicine Internal Award, Relaxera Pharmazeutische Gesellschaft to GS and NIH/NHLBI R01HL176595 and R56HL168657 to CSH.

